# Context-induced renewal of passive but not active coping behaviours in the shock-probe defensive burying task

**DOI:** 10.1101/2022.11.17.516912

**Authors:** Alexa Brown, Melissa Martins, Isabelle Richard, Nadia Chaudhri

**Affiliations:** Center for Studies in Behavioral Neurobiology, Department of Psychology, Concordia University, Montreal, QC, Canada

**Keywords:** Renewal, context, passive avoidance, active avoidance, coping responses

## Abstract

Renewal is the return of extinguished responding after removal from the extinction context. Renewal has been extensively studied using classical aversive conditioning procedures that measure a passive freezing response to an aversive conditioned stimulus. However, coping responses to aversive stimuli are complex and can be reflected in passive and active behaviours. Using the shock-probe defensive burying task, we investigated whether different coping responses are susceptible to renewal. During conditioning, male, Long-Evans rats were placed into a specific context (context A) where an electrified shock-probe delivered a 3 mA shock upon contact. During extinction, the shock-probe was unarmed in either the same (context A) or a different context (context B). Renewal of conditioned responses was assessed in the conditioning context (ABA) or in a novel context (ABC or AAB). Renewal of passive coping responses, indicated by an increased latency and a decreased duration of shock-probe contacts, was observed in all groups. However, renewal of passive coping, measured by increased time spent on the side of the chamber opposite the shock-probe, was only found in the ABA group. Renewal of active coping responses linked to defensive burying was not observed in any group. The present findings highlight the presence of multiple psychological processes underlying even basic forms of aversive conditioning and demonstrate the importance of assessing a broader set of behaviours to tease apart these different underlying mechanisms. The current findings suggest that passive coping responses may be more reliable indicators for assessing renewal than active coping behaviours associated with defensive burying.

## Introduction

In Pavlovian conditioning, extinction refers to the progressive decline in responding to a conditioned stimulus (CS) that is no longer followed by an unconditioned stimulus (US). Extinction does not erase the original CS-US association, but rather results in new inhibitory learning (Pavlov, 1927). The inhibitory learning that occurs in extinction is thought to be context-specific, since a change in context after extinction can trigger a return of extinguished conditioned responding, termed ‘renewal’ (Bouton & Bolles, 1979; Bouton & King, 1983; Bouton et al., 2006). Context-dependent renewal can be observed in ABA, ABC, and AAB designs, in which consecutive letters refer to the different contextual configurations present during conditioning, extinction, and the renewal test (Bouton & Bolles, 1979; Bouton & Ricker, 1994). Renewal is most robust in the ABA design, in which an increase in conditioned responding occurs upon a return to the original conditioning context (context A) following extinction in a different context (context B) (Bouton & Bolles, 1979; Bouton & King, 1983; Bouton & Peck, 1989). However, renewal can also occur when conditioning, extinction, and renewal test are all conducted in distinct contexts (ABC renewal; Bouton & Bolles, 1979; Bouton & Brooks, 1993), or when conditioning and extinction are conducted in the same context and the renewal test is conducted in a different context (AAB renewal; Bouton & Ricker, 1994).

Renewal of aversive conditioned responses has predominantly been investigated using aversive Pavlovian conditioning procedures in which a CS (e.g., tone) is paired with an aversive US (e.g., footshock), and passive freezing in response to the CS is used as a measure of aversive conditioning (Maren, 2001; Corcoran & Maren, 2001; Hobin et al., 2003, 2006; Knapska & Maren, 2009). The conditioned suppression task has also been used, in which animals first learn to perform an operant response (e.g., lever-press) for appetitive reinforcement (e.g., food pellet), and then a CS (e.g., tone) is paired with an aversive US (e.g., footshock). Animals learn to suppress ongoing behaviour in response to the CS and this results in a decrement of the food-seeking response, which is used as a measure of aversive conditioning (Bouton & Bolles, 1979; Bouton & Ricker, 1994). In both procedures, the CS-footshock association is extinguished by repeatedly presenting the CS in the absence of shock, which causes responding to diminish. During the renewal test, animals are either returned to the training context (ABA) or removed from the extinction context (ABC and AAB), which triggers the return of conditioned responding to the CS. Similar renewal effects have been observed when comparing ABA renewal with either ABC or AAB renewal (Corcoran & Maren, 2004), although others have found ABC and AAB renewal to be substantially weaker than ABA renewal (Bouton & King, 1983; Tamai & Nakajima, 2000; see also Zironi et al., 2006; Bossert et al., 2004; Crombag & Shaham, 2002; Nakajima et al., 2000; Khoo et al., 2020, for failures to detect ABC or AAB renewal in other conditioning preparations).

Compared to passive avoidant behaviours such as freezing or conditioned suppression, less is known about the contextual control of active avoidant behaviours. In a typical active avoidance paradigm, a Pavlovian CS (e.g., tone) is paired with an aversive US (e.g., shock) and rodents can perform an operant response to actively avoid shock (e.g., moving to the opposite side of a shuttle box, pressing a lever, or stepping onto an elevated platform). Active avoidance learning is thought to be due to a combination of Pavlovian conditioning to the CS that predicts the aversive outcome, and operant conditioning in which the operant response avoids the potential threat (Manning et al., 2021). Avoidance tasks have been used to investigate the neural underpinnings of active avoidance learning and extinction (for a review see Moscarello & Penzo, 2022), the procedural variables that influence the expression of active avoidance (Galatzer-Levy et al., 2014; Fanselow et al., 2019), and potential sex differences in the expression of active avoidance (Gruene et al., 2015; Totty et al., 2021; Mitchell et al., 2022). However, few studies have investigated the role of contextual processing in the extinction of active avoidance, and to the best of our knowledge, only one study has investigated context-induced renewal of active avoidance (Nakajima, 2014). Using a shuttle box, rats in this study were trained to move to the opposite side upon presentation of a shock-predictive CS in context A. Rats then underwent extinction either in the same context or in a different context B, and ABA, ABC and AAB renewal of active avoidance was assessed. All three forms of renewal were found to be similar in magnitude, suggesting that extinction of active avoidance is highly context-specific since a change from the extinction context elicits robust renewal (Nakajima, 2014).

An alternative animal model of aversive conditioning is the shock-probe defensive burying (SPDB) task (Pinel & Treit, 1978). In this task, rats freely explore a behavioural chamber in which an electrified shock-probe mounted on one of the walls above the surface of the bedding delivers an electric shock upon contact. The SPDB task elicits a number of quantifiable behaviours. These behaviours can be viewed as indicators of the mental state of fear/anxiety, and have traditionally been categorized as either passive or active coping responses to aversive stimuli (De Boer & Koolhaas, 2003). In the SPDB task, passive coping strategies can include spending more time away from the shock-probe (passive avoidance), making fewer contacts with the shock-probe, and remaining immobile (freezing). Rats also display active coping behaviours (Pinel & Treit, 1978; De Boer & Koolhaas, 2003; Fucich & Morilak, 2018) including ‘defensive burying’ in the SPDB task, in which the shock-probe is buried by pushing bedding from the floor forward using thrusting movements with the forepaws or snout (Pinel et al., 1980). Rats also use defensive burying for other aversive stimuli including flash bulbs, plastic tubing that delivers a burst of air, and mousetraps (Terlecki et al., 1979). Furthermore, rats bury a shock-probe that is armed more than a control probe that is unarmed, indicating that defensive burying is a learnt behaviour (Pinel & Treit, 1978; Pinel et al., 1980). During extinction, when the shock-probe is no longer electrified, rats cease to bury the probe (Pinel et al., 1985). Therefore, the SPDB task can be used to study differences in the acquisition, extinction, and renewal of both passive and active coping behaviours in response to an aversive stimulus.

The SPDB task is thought to be a more ethological model of aversive learning than traditional aversive Pavlovian conditioning procedures in which the shock administered by the experimenter is unavoidable (Treit, Pinel & Fibiger, 1981; Rodgers et al., 1997), and the observation of conditioned responses is usually limited to freezing (Gruene et al., 2015). The SPDB task is also not dependent on appetitive motivation, since it does not involve food or drug reinforcement which are present in some aversive conditioning procedures involving approach-avoidance conflict or conditioned suppression (Treit, Pinel & Fibiger, 1981; Treit & Pesold, 1990). Importantly, the SPDB task allows the aversive stimulus to be actively or passively avoided as seen in naturalistic environments (Owings & Coss, 1977), and defensive burying may be analogous to the active burying of entrance holes to underground burrows that wild rats use to avoid predators (De Boer & Koolhaas, 2003). The relative use of passive and active defensive strategies is thought to be determined by the current level of threat imminence (Fanselow & Lester, 1988; Perusini & Fanselow, 2015). During high levels of imminent threat, rapid species-specific circa-strike reactions, such as vigorous escape attempts and darting, can provide an effective response (Fanselow et al., 2019; Gruene et al., 2015). Defensive burying is considered an active avoidant behaviour that may be expressed when threat imminence is high (De Boer & Koolhaaus, 2003). When threat is less imminent, behaviour is more flexible and includes passive freezing to avoid detection. While passive defensive states linked to low threat imminence may be associated with the mental state of “fear”, active defensive states linked to high threat imminence may be associated with “panic” (Fanselow et al., 2019).

Although the SPDB task allows a more ethological spectrum of passive and active coping responses to be observed, renewal has not been investigated using the SPDB task, and the capacity of different coping responses to return after extinction is unknown. In the present experiment, we used the SPDB task to investigate ABA, ABC and AAB renewal of active and passive coping responses. During conditioning, rats were placed in a distinct context, consisting of tactile, visual, olfactory, and auditory stimuli (context A), and the shock-probe delivered a constant 3-mA shock upon contact. During extinction, rats were either placed in the same context used in training (context A), or in a different context (context B), but contacts with the shock-probe did not result in shock. Renewal in the ABA group was tested in the original training context (context A), and renewal in the ABC group was tested in a novel context (context C). Finally, the AAB group underwent extinction in the same context as training, and renewal was tested in a different context (context B). We predicted the strongest renewal of both active and passive coping responses in the ABA design, and a stronger renewal of passive versus active coping responses in each of the three groups.

## Methods Animals

Subjects were 37 experimentally naïve, male Long-Evans rats (Charles River, St. Constant, Quebec, Canada; 220-240 g upon arrival). Rats were maintained in a climate-controlled (21°C) room on a 12-h light/dark cycle with lights turned on at 7:00 h. All procedures occurred during the light phase. Rats were individually housed in standard cages (44.5 cm × 25.8 cm × 21.7 cm) containing beta chip bedding (Aspen Sani chips; Envigo, Indianapolis IN) with unrestricted access to water and food (Agribands, Charles River). Each cage contained a nylabone toy (Nylabones; Bio-Serv, Flemington, NJ), a polycarbonate tunnel (Rat Retreats, Bio-Serv) and shredded paper for enrichment.

During a nine-day acclimation period to the colony room, rats were handled, and body weight was recorded daily. Following acclimation, rats were assigned into one of three groups: ABA, ABC and AAB matched according to body weight. Rats were excluded for failure to extinguish if they spent >65 % of the session length avoiding the side of the chamber containing the shock-probe during the final extinction session (ABA, n=1; ABC, n=1; AAB, n=2). The final number of rats in each group was 11. All procedures followed the guidelines of the Canadian Council for Animal Care and were approved by the Concordia University Animal Research Ethics Committee.

### Apparatus

The SPDB task, adapted from Pinel and Treit (1978), was conducted using three identical Plexiglas chambers (40 cm × 30 cm × 40 cm). Each chamber was encased in a sound-attenuating melamine cubicle. On the left sidewall of the chamber, 5 cm above the floor of the chamber, and 1 cm above the surface of the bedding, was a hole through which the removable shock-probe could be inserted. The shock-probe (12.7 cm in length and 0.97 cm in diameter) was constructed of ABSplus-P430 thermoplastic using a 3D printer (Stratasys Dimension uPrint). The shock-probe was wrapped with a 18AWG bare, tinned copper bus bar wire (Beldon). Two cameras were used to videotape the sessions for offline behavioural scoring. One camera was mounted to the top of the melamine cubicle pointing downwards, and the second camera was positioned 30 cm from the front of the conditioning chamber.

Three contextual configurations were used which differed in visual, tactile, olfactory, and auditory elements and were in different laboratory rooms. Context 1 consisted of 0.5 inch black and yellow vertical striped walls, ¼ inch corncob bedding (7097 Teklad; Envigo, Madison, WI), lemon odor, and no background noise. Context 2 consisted of white walls with a large black star centered on each wall, dry cellulose bedding (7070C Teklad Diamond; Envigo), almond odor, and a fan within the melamine cubicle. Context 3 consisted of white walls with small red polka-dots (2 cm in diameter), aspen woodchip bedding (7093 Teklad, Envigo), cedar wood odor, and a fan within the laboratory room. Odors were prepared by diluting lemon oil (W262528-1, Sigma Aldrich), benzaldehyde (B1000, ACP Chemicals) or cedar wood oil (8000-27-9, Fisher Chemicals) with water to make a 10% solution.

### Procedure

Prior to each session, a new layer of bedding was added to the chamber floor and was smoothed to a height of 4 cm. The appropriate odor was sprayed (2.6 ml) onto the bedding. Rats were placed in the conditioning chamber on the side of the chamber opposite the shock-probe, facing away from the probe (Pinel and Treit, 1978). Prior to conditioning, all rats received three daily 10 min habituation sessions on consecutive days in the absence of the shock-probe. Rats were habituated once to each contextual configuration and the order of context habituations was counterbalanced across rats.

Rats received three daily 15 min conditioning sessions beginning 24 h after the final habituation session. The shock-probe was present during the conditioning phase and delivered a constant 3 mA shock upon contact. The context used for conditioning was counterbalanced across groups and was referred to as ‘context A’.

At 24 h after the final conditioning session, rats received five daily 15 min extinction sessions. Extinction sessions were identical to conditioning sessions except that the shock-probe was not electrified. In the AAB group, extinction occurred in the conditioning context (context A). In the ABA and ABC groups, extinction occurred in a different context from conditioning (context B).

At 24 h after the final extinction session, rats received two counterbalanced test sessions on consecutive days, in which the shock-probe was not electrified. A control test was conducted on one day, in which rats were tested in the same context as extinction, and the renewal test was conducted on the other day. In the ABA group, the renewal test occurred in the conditioning context (context A), and in the ABC and AAB groups the renewal test occurred in a different context (context C and B, respectively). **Data Analysis**

Selected behavioural variables (Table 1) were scored from video recordings using Behavioral Observation Research Interactive Software (BORIS) v6.3.1 (Friard & Gamba, 2016) by two experimenters who were blind to the experimental conditions.

**Table 1.** Behavioural variables in the shock-probe defensive burying task and associated presumed psychological processes (De Boer & Koolhaas, 2003).

| Variable (measure) | Definition | Presumed psychological process |
| --- | --- | --- |
| Passive avoidance (duration) | Time spent on the side of the chamber opposite the shock-probe in the absence of freezing | Passive coping |
| Freezing/immobility (duration) | The absence of any movements, excluding those required for respiration | Passive coping |
| Shock-probe contact (frequency, duration, latency) | Direct contact made with the shock-probe | Passive coping |
| Defensive burying (duration, latency) | Burying the shock-probe using available bedding from the chamber floor | Active coping |
| Height of bedding (cm) | Height of the peak of accumulated bedding surrounding the shock-probe | Active coping |
| Shock reactivity (intensity scale) | Reactivity to shock measured using a 4-point scale (*1=flinch involving head or paw, 2=whole-body flinch with or without slow ambulation away from probe, 3=whole-body flinch or jumping followed by immediate ambulation away from probe, 4=whole-body flinch or jumping, accompanied by the rat running to the opposite end of the chamber*) (Shah and Treit, 2004). | Reactivity |

Inter-rater reliability between the two coders was assessed for all behavioural variables (r=.88 -.92). The acquisition and extinction of conditioned responding was assessed separately using mixed analyses of variance (ANOVA) with group (ABA, ABC or AAB) and session as factors. Renewal of conditioned responding was assessed using a mixed ANOVA with group (ABA, ABC or AAB) and context (renewal or extinction) as factors. Greenhouse-Geisser corrections are reported following violations of Mauchly’s test of sphericity. Post-hoc analyses were corrected for multiple comparisons using the Bonferroni adjustment. All data analyses were conducted using IBM SPSS v21.0 (IBM Corp., Armonk, NY). Results were considered statistically significant at p<.05.

## Results

### Number of Shocks and Shock Reactivity

When the shock-probe was electrified during conditioning, all groups received a similar number of shocks (Fig. 1A; Group, 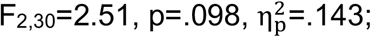 Group x Session, 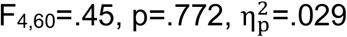) and the number of shocks decreased significantly across sessions (Session, 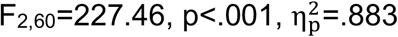). Reactivity to shock was measured using a 4-point scale (1=flinch involving head or paw, 2=whole-body flinch with or without slow ambulation away from probe, 3=whole-body flinch or jumping followed by immediate ambulation away from probe, 4=whole-body flinch or jumping, accompanied by the rat running to the opposite end of the chamber) (Shah and Treit, 2004). Groups showed comparable reactivity to shocks during acquisition (Fig. 1B; Group, 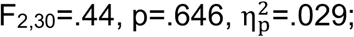 Group x Session, 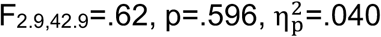), and reactivity decreased significantly across conditioning sessions (Session, 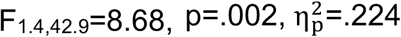).

**Fig. 1.**
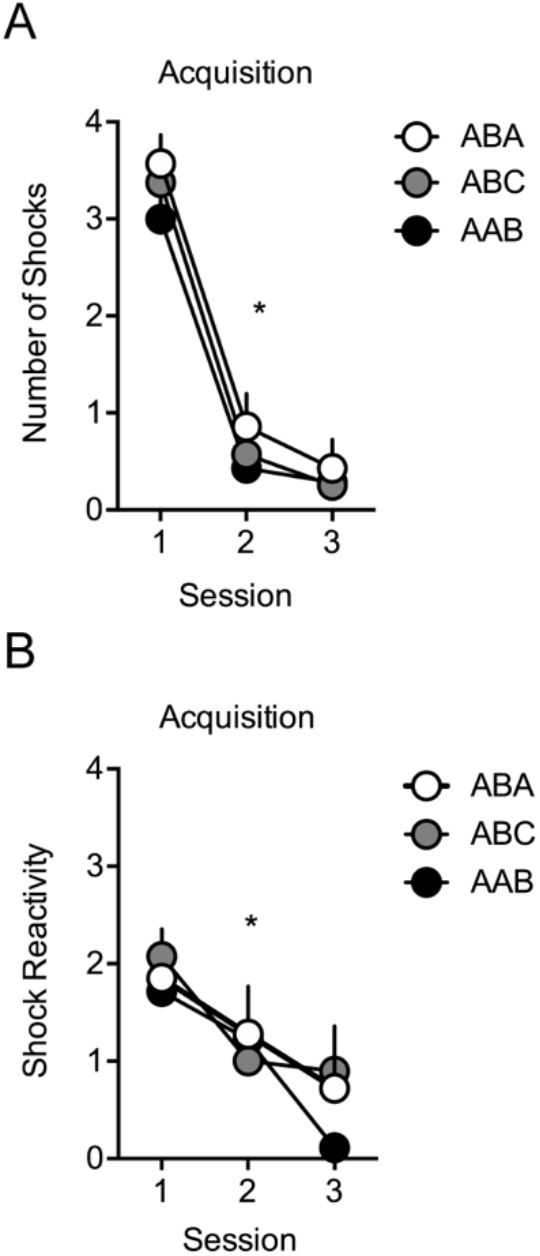
Average number of shocks and shock reactivity during acquisition in the ABA, ABC and AAB groups. **A** Average number of shocks received during the three conditioning sessions of the acquisition phase. **B** Average shock reactivity scores during the three conditioning sessions. * p<.05, main effect of session (A, B). Error bars indicate the standard error. n=11 per group.

After reacting to contacting the armed shock-probe in context A, animals in all groups displayed freezing/immobility. The duration of freezing was low in the first conditioning session (ABA, 6.4 ± 2.9 s; ABC, 2.3 ±.8 s; AAB, 2.6 ±.9 s), and decreased across sessions similarly in all groups (Data not shown; Session, 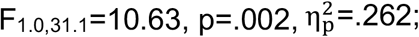 Group, 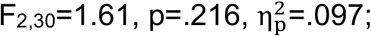 Group x Session, 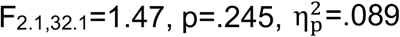). Freezing was not observed during extinction or during renewal and extinction tests when the shock-probe was unarmed.

### Passive Avoidance

#### Acquisition and Extinction

Passive avoidance was measured as time spent on the side of the chamber opposite of the shock-probe. Passive avoidance during conditioning was high across groups (Fig. 2A; Group, 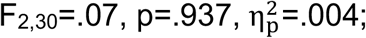 Session, 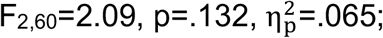 Group x Session, 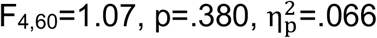), and decreased similarly across groups during extinction (Fig. 2A; Session, 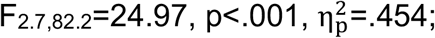 Group, 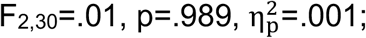 Group x Session, 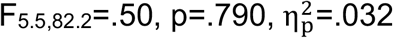). Time spent on both sides of the chamber was roughly equivalent at the end of extinction. Therefore, rats in all groups learned to passively avoid the shock-probe during conditioning, and extinguished passive avoidance during extinction.

**Fig. 2.**
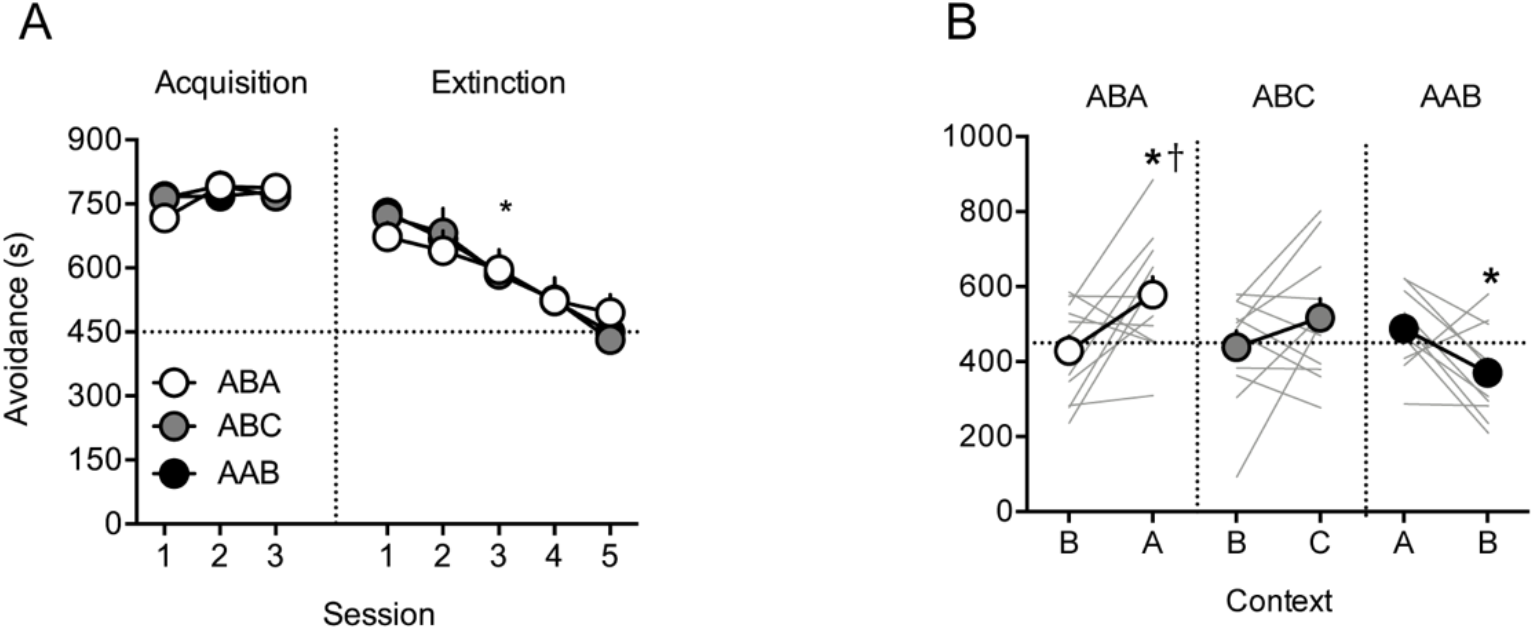
ABA renewal, but no ABC, and reverse AAB renewal of passive avoidance in rats. The horizontal dotted line represents half of the session length. **A** Acquisition and extinction of avoidance in the ABA, ABC and AAB groups. Data represent time in seconds on the side of the chamber that did not contain the shock-probe. * p<.05, main effect of session. Error bars indicate the standard error. **B** Each rat was tested once in the extinction context and once in the renewal context. Passive avoidance was significantly increased in the renewal context compared to the extinction context in the ABA group and was significantly decreased in the renewal context compared to the extinction context in the AAB group. In the ABC group, there was no significant difference between the extinction and renewal contexts. * p<.05, extinction vs. renewal context. † p<.05, ABA vs. AAB in the renewal context. Error bars indicate the standard error and data shown for each individual rat are overlaid on the graph. n=11 per group.

#### Renewal Test

Renewal of passive avoidance was observed in the ABA group, but not in the ABC or AAB groups (Fig. 2B; Group, 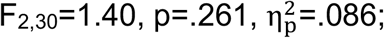 Context, 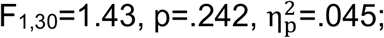 Group x Context, 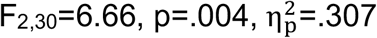). Passive avoidance was significantly increased in the renewal context compared to the extinction context in the ABA group 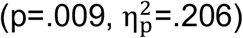 but did not reach statistical significance in the ABC group 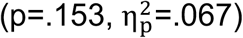. In contrast, passive avoidance was significantly reduced in the renewal context compared to the extinction context in the AAB group 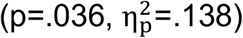. Therefore, we observed renewal of passive avoidance in the ABA group, and a suppression of avoidance in the AAB group.

### Shock-Probe Contact

#### Acquisition and Extinction

Measures of shock-probe contact showed similar changes during conditioning in all groups, and during the five extinction sessions. During acquisition, the frequency of shock-probe contacts decreased significantly in all groups (Fig. 3A; Session, 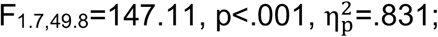 Group, 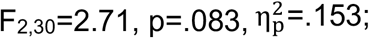 Group x Session, 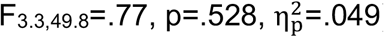), the duration of shock-probe contacts was reduced (Fig. 3B; Session, 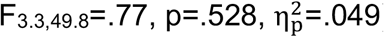 Group, 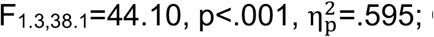 Group x Session, 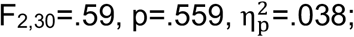) and the latency to contact the shock-probe was increased (Fig. 3C; Session, 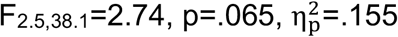 Group, 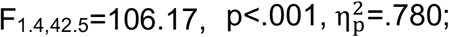 Group x Session, 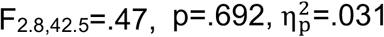).

**Fig. 3.**
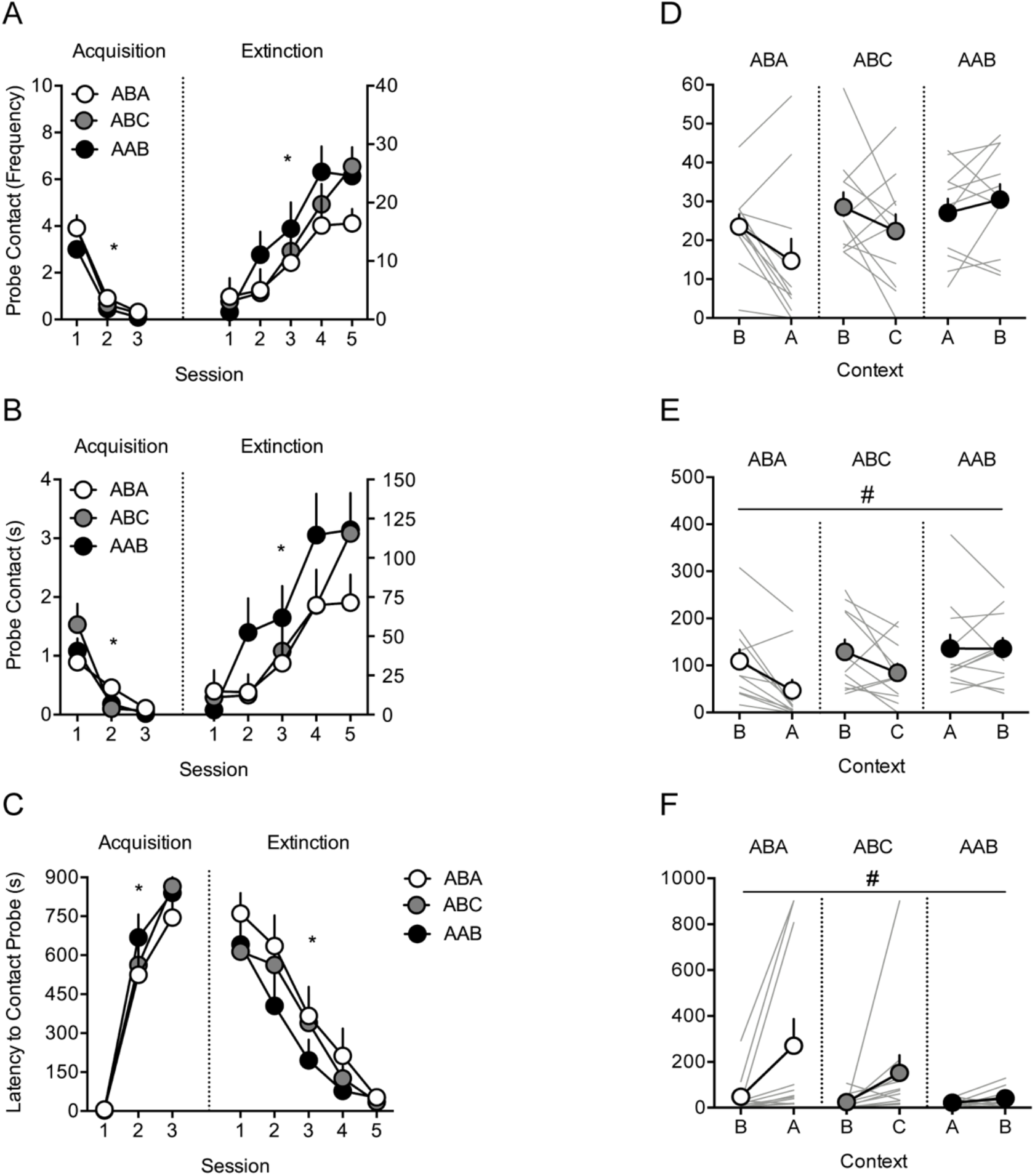
Renewal of shock-probe contact duration and latency across groups, but no renewal of shock-probe contact frequency. **A** Average frequency of shock-probe contacts, **B** Average duration of shock-probe contacts, and **C** Average latency to initiate shock-probe contact during the acquisition and extinction phases in the ABA, ABC and AAB groups. * p<.05, main effect of session (A, B, C). Error bars indicate the standard error. **D** Each rat was tested once in the extinction context and once in the renewal context. There was no difference between extinction and renewal contexts for the frequency of shock-probe contacts. **E** The duration of shock-probe contacts was significantly decreased in the renewal context compared to the extinction context. **F** The latency to initiate shock-probe contact was significantly increased in the renewal context compared to the extinction context. # p<.05, main effect of context (E, F). Error bars indicate the standard error and data shown for each individual rat are overlaid on the graph. n=11 per group.

Responses in all three groups were also similar during extinction, in which the frequency of shock-probe contacts was increased (Fig. 3A; Session, 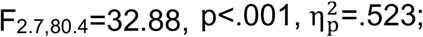 Group, 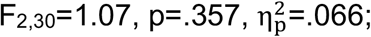 Group x Session, 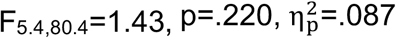), the duration of shock-probe contacts was increased (Fig. 3B; Session, 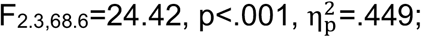 Group, 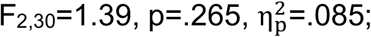 Group x Session, 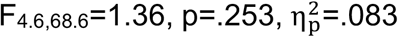) and the latency to contact the shock-probe was reduced (Fig. 3C; Session, 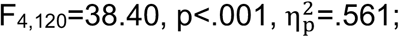 Group, 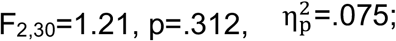 Group x Session, 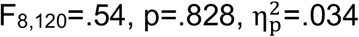). Therefore, rats across all renewal groups learned that the shock-probe was aversive and avoided contact during conditioning, and increased contact during extinction when the probe was unarmed.

#### Renewal Test

Renewal of shock-probe contact was expressed by shock-probe contact durations and latencies to contact the shock-probe. Shock-probe contact duration was significantly reduced in the renewal context compared to the extinction context across groups (Fig. 3E; Context, 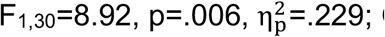 Group, 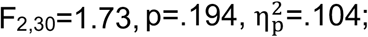 Group x Context, 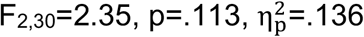), and the latency to contact the shock-probe was increased in the renewal context compared to the extinction context across groups (Fig. 3F; Context, 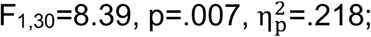 Group, 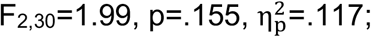 Group x Context, 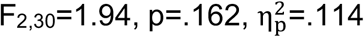). However, the frequency of shock-probe contacts did not differ significantly between the renewal context and the extinction context (Fig. 3C; Context, 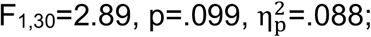 Group, 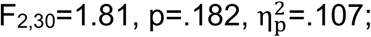 Group x Context, 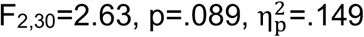). Therefore, we observed renewal of passive coping measured by shock-probe contact duration and latency, and these effects appeared to be primarily driven by the ABA group, followed by the ABC group, and weak renewal in the AAB group.

### Active Defensive Burying

#### Acquisition and Extinction

Measures of active defensive burying during conditioning showed similar changes in all groups. The duration of defensive burying decreased across sessions in all groups (Fig. 4A; Session, 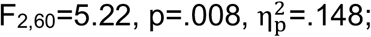 Group, 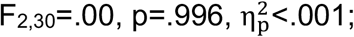 Group x Session, 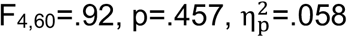). However, there were no significant changes in the latency to initiate defensive burying (Fig. 4B; Session, 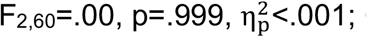 Group, 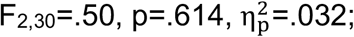 Group x Session, 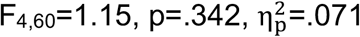) or in the height of the accumulated bedding surrounding the shock-probe (Fig. 4C; Session, 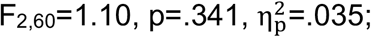 Group, 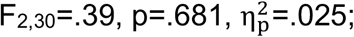 Group x Session, 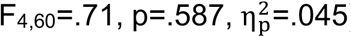).

During extinction, all groups ceased to bury the shock-probe. The duration of defensive burying decreased across sessions in all groups (Fig. 4A; Session, 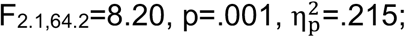 Group 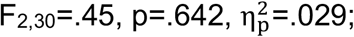 Group x Session, 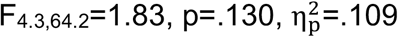), and the latency to initiate defensive burying was increased (Fig. 4B; Session, 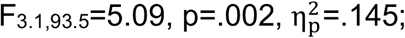 Group, 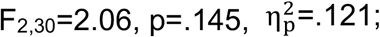 Group x Session, 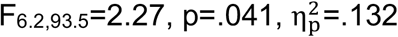). The ABC group had shorter latencies to bury during extinction session 2 compared to the ABA (p=.015, d=1.37) and AAB (p=.035, d=1.18) groups, and during extinction session 3 compared to the ABA group (p=.044, d=1.20). However, all groups reached similar latencies by the final extinction session. The height of the peak of accumulated bedding surrounding the shock-probe decreased comparably across sessions in all groups (Fig. 4C; Session, 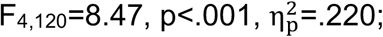 Group 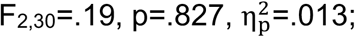 Group x Session, 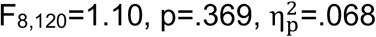). Therefore, rats in all groups showed defensive burying behaviour during conditioning and reduced defensive burying during extinction.

**Fig. 4.**
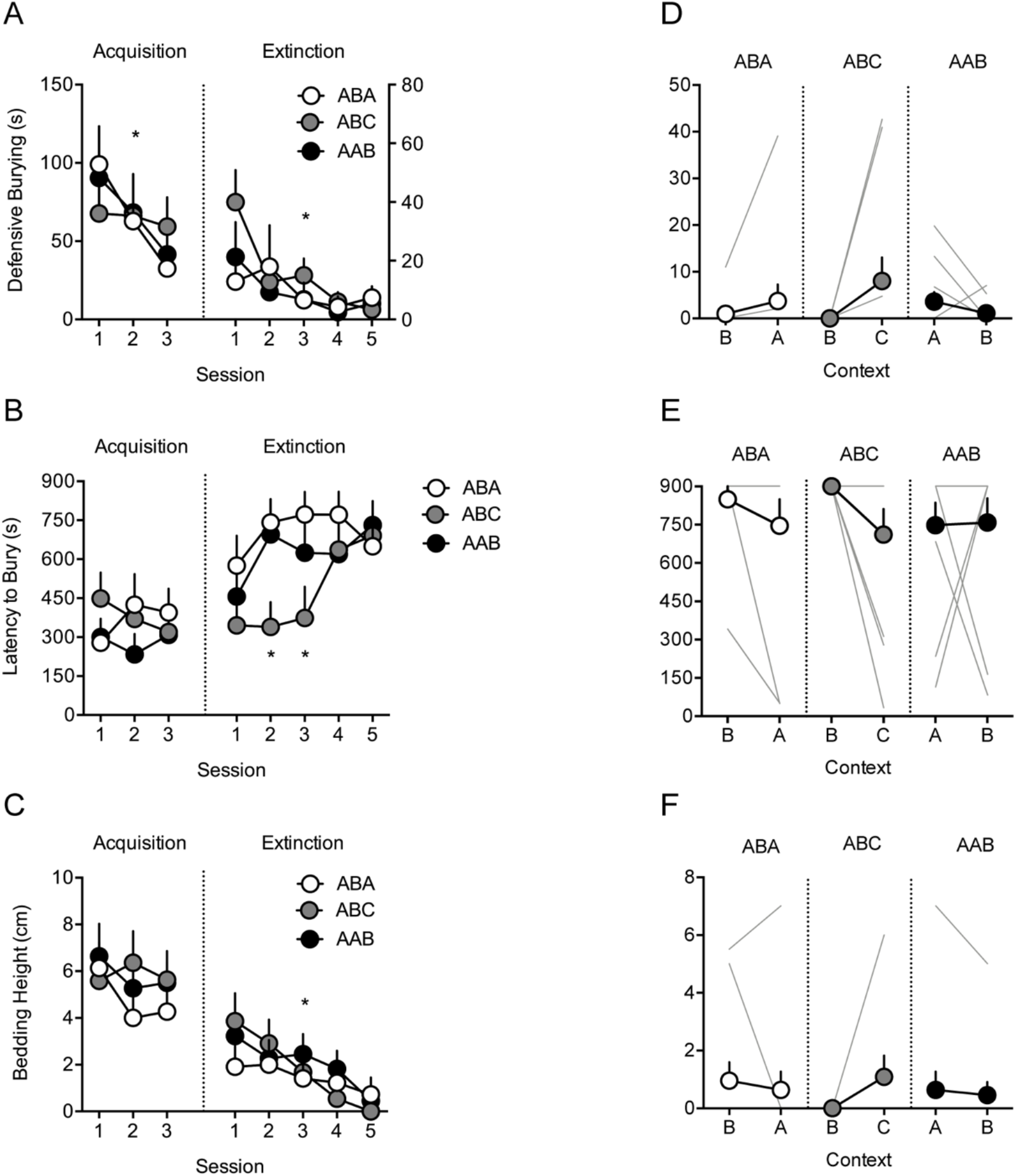
There was no renewal of active defensive burying in the ABA, ABC or AAB groups. **A** Average duration of defensive burying, **B** Average latency to initiate defensive burying, and **C** Average height of the peak of accumulated bedding material surrounding the shock-probe during the acquisition and extinction phases. * p<.05, main effect of session (A, C). *p<.05, significantly different responding in the ABC group compared to the ABA and AAB groups in extinction session 2, and significantly different responding in the ABC vs. ABA group in extinction session 3 (B). Error bars indicate the standard error. **D** Each rat was tested once in the extinction context and once in the renewal context. There was no difference between extinction and renewal contexts for the duration of defensive burying, **E** the latency to initiate defensive burying, or **F** the height of the peak of accumulated bedding material surrounding the shock-probe. Error bars indicate the standard error and data shown for each individual rat are overlaid on the graph. For rats that showed no response, individual data is at 0 (D, F) and 900 (E). n=11 per group.

#### Renewal Test

Measures of defensive burying behaviour did not show significant renewal in any group. There was no significant difference between the renewal and extinction contexts in all groups for the duration of defensive burying (Fig. 4D; Context, 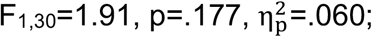 Group, 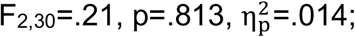 Group x Context, 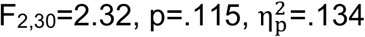), the latency to initiate defensive burying (Fig. 4E; Context, 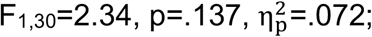 Group, 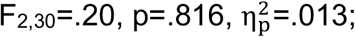 Group x Context, 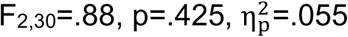), or the height of the accumulated bedding (Fig. 4F; 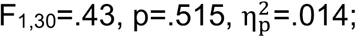 Group, 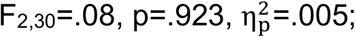 Group x Context, 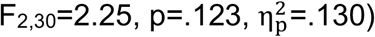). Therefore, we did not observe renewal of active coping strategies measured by duration and latency of defensive burying, and height of the peak of accumulated bedding surrounding the shock-probe.

## Discussion

The present experiment investigated ABA, ABC and AAB renewal of active and passive coping behaviours using the shock-probe defensive burying (SPDB) task in rats to determine whether these coping strategies are differentially subject to renewal.

During conditioning, rats in all three groups passively avoided the side of the chamber containing the shock-probe and reduced contact with the probe, and actively buried the shock-probe using bedding from the chamber floor. During extinction, all groups showed similar reductions in passive avoidance, an increase in shock-probe contact, and a reduction in active defensive burying. Increased passive avoidance in the renewal test was significant for the ABA group but did not reach statistical significance in the ABC group. In contrast, in the AAB group, rats spent less time passively avoiding the shock-probe in the novel context as compared to the extinction context. Moreover, at test, we detected renewal of passive coping measured by shock-probe contact duration and latency, which appeared to be primarily driven by the ABA group, and weak renewal in the ABC and AAB groups. Our observation of context-dependent renewal of passive coping strategies, in the absence of renewal of active coping behaviours associated with defensive burying, suggests that passive coping behaviours may serve as stronger indicators for examining the renewal effect in associative learning studies in comparison to defensive burying-related active coping behaviours.

### Acquisition and Extinction

The SPDB task is an ethological model of aversive conditioning that allows for the expression and measurement of multiple naturalistic behaviours (Pinel & Treit, 1978; De Boer & Koolhaas, 2003). Other aversive conditioning tasks, such as Pavlovian fear conditioning and conditioned suppression, allow assessment of only passive coping behaviours such as freezing (e.g., Bouton & Bolles, 1979; Bouton & King, 1983; Corcoran & Maren, 2001; Hobin et al., 2003). The SPDB task allows assessment of passive coping through measures of shock-probe contact and time spent avoiding the side of the chamber containing the shock-probe, and also allows assessment of active coping including the duration and latency of defensive burying, and height of the peak of accumulated bedding surrounding the shock-probe, which is analogous to natural responses to predation (De Boer & Koolhaas, 2003).

During acquisition in context A, we found that rats in the three groups similarly expressed passive and active coping behaviours. Freezing was only observed during acquisition, as a response to the shock, and was less frequent than passive avoidance. Similar levels of extinction of passive and active behaviours were observed in all three groups when the shock-probe was unarmed. Our finding that extinction was similar in the same context as conditioning (AAB) as when extinction was conducted in a different context as conditioning (ABA and ABC) is consistent with reports in Pavlovian learning where switching the extinction context after conditioning has no considerable effect on responding to the CS (Bouton & King, 1983; Bouton & Brooks, 1993). Moreover, Rosas and colleagues (2007) demonstrated that extinction of a taste aversion response is similarly expressed when extinction occurs in the conditioning context (AAB) and following a switch in context after conditioning (ABA), suggesting that across a variety of aversive learning procedures switching the extinction context after conditioning does not affect extinction. Together, the acquisition and extinction of passive and active coping responses allow for the observation of renewal outside of the extinction context.

### Renewal of passive coping strategies

Renewal involves the return of conditioned behaviours after removal from the extinction context. In studies of Pavlovian fear conditioning and conditioned suppression, renewal is expressed most strongly when animals are re-exposed to the original conditioning context (Bouton & King, 1983; Tamai & Nakajima, 2000), reflecting the importance of contextual cues for renewal (Bouton, 2004). We assessed the role of context in the SPDB task by comparing renewal in the ABA design, in which renewal should be most robust, with the ABC and AAB renewal designs in which renewal occurs in a novel context. Comparisons of passive coping behaviours in the renewal context versus the extinction context at test indicated that the ABA group showed renewal through an increase in the latency to initiate probe contact, reduced probe contact duration, and increased passive avoidance characterised by increased time spent on the side of the chamber opposite the shock-probe. This finding of robust ABA renewal of passive coping strategies is consistent with Pavlovian freezing and conditioned suppression studies (Bouton & Bolles, 1979; Bouton & King, 1983; Bouton & Peck, 1989; Corcoran & Maren, 2001, 2004; Hobin et al., 2006). Renewal of freezing behaviour was not observed, and this may be due to the low levels of freezing observed just following reactions to probe contacts during conditioning. Low levels of freezing have been observed in both ethological and experimental studies, which show that rats freeze for less than ∼10% of the session length (De Boer & Koolhaas, 2003; Tao et al., 2017).

Renewal of passive coping strategies in the SPDB task was much less marked in the ABC and AAB groups. A subset of rats in the ABC group showed a renewal of passive avoidance but there was no significant renewal in the ABC group (p=.153). Studies directly comparing ABA and ABC renewal in aversive learning procedures are sparse. ABA and ABC renewal have been shown to be similar in magnitude using a conditioned suppression task (Thomas et al., 2003) and a signalled avoidance task (Nakajima, 2014). However, in appetitive conditioning procedures, ABA renewal has been shown to be a more robust effect compared to ABC renewal (Zironi et al., 2006; Khoo et al., 2020).

AAB renewal has been observed in a variety of learning tasks (Bouton & Ricker, 1994; Rosas et al., 2007; Rescorla, 2007, 2008; Bouton et al., 2011; Todd et al., 2013), but other studies have also been unable to detect AAB renewal (Nakajima et al., 2000; Crombag & Shaham, 2002; Bossert et al., 2004; Fuchs et al., 2005; Khoo et al., 2020). Moreover, AAB renewal has been found to be much weaker than ABA and ABC renewal in fear conditioning (Thomas et al., 2003). We observed the opposite of a renewal effect in the AAB group such that passive avoidance was reduced in the renewal context compared to the extinction context. AAB renewal is not easily explained by many learning theories (e.g., Pearce, 1987; Rescorla & Wagner, 1972) and AAB suppression of conditioned responding has only been previously observed using an appetitive Pavlovian conditioning procedure (Khoo et al., 2020). Khoo and colleagues (2020) suggested that AAB suppression may be due to the AAB group experiencing extinction of the stimulus and of context A, presumably resulting in an inhibitory memory that is strong enough to prevent renewal in context B. Similarly, Laborda and colleagues (2011) have suggested that AAB renewal is a weak effect because the AAB group undergoes “deeper extinction” resulting in weaker AAB renewal compared of ABC renewal.

Although rats were habituated to contexts prior to testing, it is possible that the AAB suppression of responding in the present study may be due to the perceived novelty of context B at test. Rats have a natural tendency to explore novel environments and familiar objects in novel contexts (Mumby et al., 2002), and it is possible that the novelty of context B at test might have driven more exploratory behaviour during the AAB renewal test, and less time on the side of the chamber opposite the shock-probe.

Context novelty could therefore have played a role in inhibiting fear responses. Bouton and Ricker (1994) found AAB renewal of conditioned behaviour in appetitive and aversive conditioning tasks. However, in their experimental preparations, rats were equally exposed to both contexts (context A and context B) prior to the renewal test. In contrast, in our study, rats in the AAB group received greater exposure to context A (9 sessions in total) compared to context B (1 session in total). Therefore, while it was expected that a change in context after extinction would lead to a return of fear responses, the perceived novelty of the context could have masked this effect and influenced behaviour in the opposite direction.

Our observation of ABA, but not ABC or AAB renewal of passive avoidance, is consistent with the interpretation that context A retains a residual excitatory association with the US (i.e., shock) after extinction, which summates with the residual excitatory strength of the CS (i.e., shock-probe) at test, and results in greater ABA renewal compared to ABC and AAB renewal (Rescorla & Wagner, 1972; Polack et al., 2013; Delamater & Westbrook, 2014). Similarly, Totty et al. (2021) found that the shift from freezing to an active flight response evoked by the CS was mediated by the summation of contextual and cued fear in a Pavlovian serial-compound stimulus task. However, the findings of Nakajima (2014), showing similar renewal of active avoidance in ABA, ABC, and AAB groups suggests that an occasion-setter mechanism mediates extinction of active avoidance. Therefore, the precise role of context in mediating avoidant behaviours remains unclear, but it is possible that the summation of both context and CS associations with the US contribute to renewal of active avoidance in some paradigms, as well as the renewal of passive avoidance observed in the present study.

### Renewal of active coping strategies

Context-induced renewal of active coping strategies measured by defensive burying was not observed in any of the groups, even in the ABA group that showed renewal of passive avoidance. Several procedural variables that are known to affect the variability of defensive burying were considered prior to testing and are unlikely to have contributed to the lack of renewal of active defensive burying. We used Long-Evans rats that are known to bury more than Wistar rats (Tarte & Oberdiek, 1982) and used a chamber size that is typical in SPDB tasks (Pesold & Treit, 1992; Degroot et al., 2001; Shah & Treit, 2004; Trent & Menard, 2013; Tao et al., 2017). The intensity of the shock administered in our experiment (3 mA) is also like that used in other studies that have observed robust defensive burying (Trent & Menard, 2013; Tao et al., 2017).

The absence of renewal of defensive burying is more likely due to the variability in the degree to which rats employed the defensive burying response strategy. We observed robust defensive burying similar to burying observed in other studies (Pesold & Treit, 1992; Shah & Treit, 2004) and, on average, defensive burying represented about 10% of the conditioning session length (85.7 ± 16.0 s). Burying was highly variable between rats however and ranged from 1.3 to 299.3 s. This is in line with previous research showing a high degree of variability in defensive burying both within- and between-studies, with defensive burying representing, on average, 3 to 30% of the observation time in standard duration test (10-15 min) (De Boer & Koolhaas, 2003). The tendency of some rats to show low degrees of defensive burying may have contributed to the absence of significant renewal.

Variability in defensive burying can also be attributed to the high degree of flexibility over behaviour in the SPDB task. Active defensive states such as defensive burying are metabolically costly (McEwan et al., 2015) and may even increase the likelihood of shock if the rats approach too close to the shock-probe (De Boer & Koolhaas, 2003). In contrast, passive coping responses require little metabolic investment and animals can avoid the shock altogether (De Boer & Koolhaas, 2003). This idea is consistent with the predatory imminence continuum (Fanselow & Lester, 1988; Perusini & Fanselow, 2015), which indicates that the perceived distance from contact with a threatening stimulus determines the selection of either passive or active responses. When rats employ a passive avoidance response by spending time on the side of the chamber opposite the shock-probe, they become spatially distanced from the threat, and this may reduce active defensive burying by reducing the perceived imminence of danger. Active coping responses such as defensive burying may therefore be most actively employed after receiving the shock during conditioning when the perceived threat is imminent and animals may experience “panic”, but not after extinction in the renewal context when the threat can be passively avoided and the sense of fear is reduced (Fanselow et al., 2019). Therefore, the SPDB task provides greater behavioural flexibility than typical active avoidance tasks, which require an active response to avoid threat. The lack of a renewal effect observed here contrasts with the findings of Nakajima (2014) who demonstrated ABA, ABC, and AAB renewal of active avoidance in a signalled shuttle box task. This suggests that the contextual control of active avoidant behaviours varies depending on the task used, and that the strongest effects may be observed when effective avoidance demands an active response.

## Conclusions

Our results show renewal of passive coping behaviours, but not renewal of active defensive burying, in the SPDB task. Consistent with renewal of passive freezing in aversive Pavlovian conditioning (Bouton & King, 1983; Tamai & Nakajima, 2000; Thomas et al., 2003), renewal of passive coping was predominantly observed in the ABA group and less so for the ABC and AAB groups. We did not detect renewal of active defensive burying in any group. These results are the first to investigate differences in context-induced renewal of passive and active coping strategies using the SPDB task. Our results provide novel evidence suggesting that active coping responses linked to defensive burying may not be expressed sufficiently to allow study of context-induced renewal as compared to passive coping responses. Active coping responses are more energetically demanding, and may be less likely to be expressed when danger is perceived as less imminent following extinction, and can be avoided passively.

## Acknowledgments

This research was funded by the Natural Sciences and Engineering Research Council of Canada. N.C. was the recipient of a Chercheur-Boursier award from the Fonds de Recherche en Santé Québec and member of the Center for Studies in Behavioral Neurobiology. A.B. is supported by a doctoral scholarship from the Fonds de Recherche du Quebec en Nature et Technologies. N.C. and A.B. designed all experiments. A.B. conducted and analyzed the data for the experiment and M.M. and I.R. helped conduct the experiment. A.B. prepared the article with input from N.C. The authors would like to thank Dr. Andrew Chapman for reviewing the manuscript and providing comments, and David Munro for technical support.

## Declarations

### Ethics approval

Methods used in this study were approved by the Concordia University Animal Research Ethics Committee.

### Consent to participate

Not applicable

### Consent for publication

Not applicable

### Code availability

Not applicable

### Authors’ contributions

N.C. and A.B. designed all experiments. A.B. conducted and analyzed the data for the experiment and M.M. and I.R. helped conduct the experiment. A.B. prepared the article with input from N.C.

### Availability of data and materials

The datasets generated for this study are available on request to the corresponding author. None of the experiments was preregistered.

